# Development of an RT-RPA assay for La Crosse virus detection provides insights into age-dependent neuroinvasion in mice

**DOI:** 10.1101/2025.01.21.634131

**Authors:** Lily Lumkong, Reem Alatrash, Sainetra Sridhar, Prince Baffour Tonto, Bobby Brooke Herrera

**Affiliations:** Rutgers Global Health Institute, Rutgers University, New Brunswick, NJ, USA; Department of Medicine, Division of Allergy, Immunology, and Infectious Diseases, and Child Health Institute of New Jersey, Rutgers Robert Wood Johnson Medical School, Rutgers University, New Brunswick, NJ, USA

**Keywords:** La Crosse virus, Reverse transcription recombinase polymerase amplification, Lateral flow assay, Neuroinvasion, Age-susceptible mouse model

## Abstract

**Background:** La Crosse virus (LACV) is a mosquito-borne arbovirus responsible for pediatric encephalitis in North America, predominantly affecting children under 16 years. Early and accurate diagnosis is critical to reducing morbidity in this vulnerable population. However, existing molecular and serological methods are limited in sensitivity, specificity, and accessibility.

**Methods:** To address these limitations, we developed a reverse transcription recombinase polymerase amplification (RT-RPA) assay for LACV detection. Primers targeting the divergent M segment of the LACV genome were designed and screened for optimal performance. The assay’s analytical sensitivity was evaluated through serial dilutions of LACV RNA prior to reverse transcription, while specificity was assessed using reverse transcribed RNA from related or geographically relevant arboviruses. We further adapted the RT-RPA test into a lateral flow assay (LFA) format for potential point-of-care use. Additionally, we employed a murine model to explore the age-dependent dynamics of LACV neuroinvasion and clearance, with the virus detected using RT-RPA and reverse transcription quantitative polymerase change reaction (RT-qPCR).

**Results:** Primer screening identified an optimal primer pair that amplified LACV cDNA within 20 minutes at 39°C, with a limit of detection (LOD) of 100 copies. The assay demonstrated high specificity, with no amplification of related or other geographically relevant arboviruses. Integration of the RT-RPA test into an LFA format preserved the LOD and specificity, enabling visual detection via test strips. In the murine model, weanling mice exhibited LACV neuroinvasion as early as 4 days post-infection (dpi), with sustained detection between 5-7 dpi. In adult mice, neuroinvasion was first detected at 5 dpi, plateauing between 6-10 dpi, and cleared entirely by 20 dpi in surviving animals.

**Conclusions:** This study establishes the RT-RPA assay as an efficient, specific, and sensitive diagnostic platform for LACV, with potential for adaptation into field-deployable LFA tests. Moreover, our findings provide valuable insights into the age-dependent dynamics of LACV neuroinvasion and clearance, informing future diagnostic and therapeutic strategies.

## Background

La Crosse virus (LACV) is a single-stranded, negative-sense virus within the *Peribunyaviridae* family, order *Bunyavirales*, and is classified under the California serogroup (CSG) viruses (1, 2). Members of this viral family possess a distinctive tri-segmented genome composing the small (S), medium (M), and large (L) segments. The S segment of LACV encodes both the nucleocapsid (N) protein and a nonstructural protein (NSs) implicated in the disruption of host innate immune responses (3, 4). The M segment encodes glycoproteins Gn and Gc, which mediate viral attachment and entry, alongside a nonstructural protein (NSm) crucial for virion assembly (5). The L segment encodes the RNA-dependent RNA polymerase (RdRp), an enzyme integral to viral genome transcription and replication (6–8).

LACV was first identified in La Crosse, Wisconsin in the 1960s, and has since remained endemic in North America, particularly in the Midwestern and Eastern United States, including the Appalachian corridor (9). Its principal vector is the Eastern treehole mosquito, *Aedes triseriatus*, although more recent data indicate that *Aedes albopictus* and *Aedes japonicus* may also contribute to transmission, raising concerns about geographic expansion (10–12). Between 2003 and 2023, the Centers for Disease Control and Prevention (CDC) documented 1,266 LACV-related hospitalizations in the United States – most of which affected pediatric patients – yet the actual incidence is likely underestimated due to underreporting and diagnostic limitations (7, 13, 14). Furthermore, snowshoe hare virus (SSHV) – a closely related CSG virus that shares approximately 89% sequence identity with LACV – circulates endemically in Canada and has also been associated with outbreaks in young populations, highlighting that arboviruses within the CSG pose a broad threat to populations across North America (15, 16).

Human infections with LACV exhibit a wide spectrum of outcomes. While most cases remain asymptomatic or present as mild febrile illness, a subset of infections progress to severe central nervous system (CNS) involvement, resulting in encephalitis, seizures, altered mental status, and, in rare instances, coma or fatality (17). Children under 16 years of age are particularly susceptible to adverse outcomes, a pattern attributed in part to differences in immune responses and blood brain barrier function (10, 18–23).

Despite its clinical importance, LACV remains difficult to diagnose. Conventional methods such as reverse transcription quantitative polymerase chain reaction (RT-qPCR) provide high sensitivity and specificity during peak viremia, yet its utility diminishes once viral titers wane (13, 24–27). Serological tools, including enzyme-linked immunosorbent assays (ELISAs), can identify LACV-specific antibodies, but often require confirmatory plaque reduction neutralization tests (PRNTs) to differentiate infections from LACV and closely related viruses (28). Moreover, the absence of commercially available LACV-specific RT-qPCRs and ELISAs, and the labor-intensive nature of PRNTs, limit the scalability of these approaches, especially in areas with restricted diagnostic resources.

To address these diagnostic hurdles, recombinase polymerase amplification (RPA) has emerged as a promising isothermal nucleic acid amplification technology (29). By functioning at 37-42°C and completing amplification in as few as 20 minutes, RPA reduces the need for specialized equipment. This method employs recombinase enzymes to bind primers to homologous sequences, single-stranded DNA-binding proteins to stabilize the resulting complexes, and strand-displacing polymerases to drive rapid amplification (30–32). When combined with reverse transcription (RT-RPA), the method allows rapid detection of RNA viruses like LACV through an initial conversion of RNA into cDNA. Furthermore, RPA amplicons can be evaluated using standard gel electrophoresis or via lateral flow assays (LFAs), the latter providing near-instantaneous visual results suitable for field-based or point-of-care settings (27, 33). LFAs capitalize on capillary action within a nitrocellulose strip, where labeled detection reagents bind to specific target amplicons, generating visible test and control lines (34–36).

In addition to the critical need for improved diagnostics, LACV exhibits age-dependent pathogenicity that requires further investigation. Notably, the age-dependent disease severity observed in humans is recapitulated in murine models, where weanling (3 week old) mice display higher mortality rates and more pronounced CNS pathology between 5-7 days post-infection (dpi) compared to adult (≥8 week old) mice with minimal to no death (7, 9, 21, 37, 38). While the pathogenesis of LACV has been relatively well studied in mice and other mammals, limited data are available regarding the temporal dynamics of LACV neuroinvasion into the CNS and/or brain (15, 39, 40).

In this study, we describe the development and validation of an RT-RPA assay to detect LACV with high sensitivity and specificity. We also utilize an age-susceptible mouse model to characterize the temporal progression of LACV infection in the brain, exploring how host age influences both neuroinvasion and viral clearance. Our study seeks to enhance early detection in endemic regions and improve the broader understanding of host-pathogen interactions in LACV encephalitis.

## Methods

### Multiple sequence alignments and primer/probe design

Percent identity matrix analysis was performed on the S, M, and L segments from a panel of CSG viruses. These included LACV (GenBank accession number: EF485031.1), SSHV (GenBank accession number: EU262553.1), Khatanga virus (GenBank accession number: KT288294.1), California Encephalitis virus (GenBank accession number: KX817313.1), San Angelo virus (GenBank accession number: KX817331.1), Lumbo virus (GenBank accession number: KX817325.1), Tahnya virus (GenBank accession number: EU622819.2), and Jamestown Canyon virus (GenBank accession number: HM007351.1). Multiple sequence alignments and homology analyses were generated using Clustal Omega (version 1.2.4). The resulting sequence alignments were subjected to phylogenetic analysis by constructing maximum-likelihood trees based on the Tamura-Nei model and 500 bootstrapping replications using Mega11 software.

Primers were manually designed using SnapGene software (version 8.0.1) (Supplementary Table 1), adhering to established RPA guidelines that limit the primer length to 30-35 bases, maintain 40-60% GC content, and minimize palindromic or repetitive sequences (32). Primer candidates were initially tested in silico for hairpin and dimer formation. Those meeting these criteria underwent validation at three temperatures, including 37°C, 39°C, and 42°C, for 20 minutes. The primer pair that yielded the strongest, most specific band was used in subsequent experiments.

### RNA Extraction

The PureLink Viral RNA/DNA Mini kit (ThermoFisher) was used for RNA extraction from LACV (BEI Resources: NR-540, GenBank accession number: GCA_004789695.1), SSHV (BEI Resources: NR-535, GenBank accession number: GCA_004789895.1), DENV-1 (BEI Resources: Hawaii strain, GenBank accession number: KM204119.1), DENV-2 (BEI Resources: DakArA1247 strain, GenBank accession number: EF105383), DENV-3 (BEI Resources: DENV-3/US/BID-V1043/2006 strain, GenBank accession number: EU482555), DENV-4 (BEI Resources: H241 strain, GenBank accession number: AY947539), ZIKV (BEI Resources: DAK AR 41524 strain, GenBank accession number: KX601166), WNV (BEI Resources: Bird114 strain, GenBank accession number: GU827998.1), and YFV (BEI Resources: YFV17D strain, GenBank accession number: MG051217.1), following the manufacturer’s instructions. Additionally, RNA from mouse brains was extracted using the RNeasy Lipid tissue Mini Kit (Qiagen), following the manufacturer’s instructions. The concentration of the RNA extracts was determined using a NanoDrop One device (ThermoFisher).

### cDNA synthesis and primer screening

Viral RNA was converted to cDNA using the High-Capacity cDNA Reverse Transcription Kit (ThermFisher), following the manufacturer’s instructions. 10 μl of extracted RNA was input into a 20 µL cDNA synthesis reaction. For the primer screen, 1 μl of viral cDNA was used as input for primer screens, limit of detection (LOD) test, and cross-reactivity screen RPA reactions. 1 μl of water was used as input for negative controls.

### RT-RPA reactions using unlabeled primers

RPA reactions were conducted using the TwistAmp Basic Lyophilized Kit (TwistDx), following the manufacturer’s instructions, but were scaled down from a 50 µL reaction to a 10 µL reaction. In brief, the lyophilized mix was rehydrated using 29.5 µL of the provided rehydration buffer and 10 µL of nuclease-free water. 7.9 µL of the rehydrated mix was used per 10 µL reaction, along with 0.3 µM of each primer, 280 mM magnesium acetate, and 1µL of cDNA. The reactions were incubated for 20 minutes at either 37°C, 39°C, or 42°C. Diluted reaction products (2:1) were run on a 2.5% agarose gel (BioRad) in 1x TAE buffer (BioRad) for 30 minutes at 100 volts.

### RT-RPA reactions using labelled primers

For the hybridization method, reactions were set up as described as above, with one minor change, where the reverse primer was replaced with a reverse primer with a 5’biotin labelled version, with the same primer concentration in the reaction. The hybridization step was performed as described by Qian’s hybridization protocol (41), using a probe that was labeled with a 5’-FAM tag (Supplementary Table 1). Following the RPA reaction run, 1 µL of 5 µM hybridization probe (final concentration of 0.167 µM) and 19 µL of hybridization buffer (10 mM Tris HCl at pH 8.5) were added to each reaction. This reaction mix was heated to 95°C for 3 minutes on the thermocycler and then cooled to room temperature to allow for the hybridization reaction.

### Detection of RPA products on LFA strips

The products from the hybridization reactions were processed using the HybriDetect 2T LFA strips (Milenia Biosciences), following the manufacturer’s instructions. Unpurified reaction products were diluted in the provided assay buffer (1:15 for hybridization reactions) and applied to the strip. The strip was then immersed in 70 µL of the provided assay buffer and was allowed to react for 10-15 minutes. The strips were imaged using a smartphone camera.

### qPCR

cDNA samples derived from mice brains were analyzed by qPCR using PowerTrack SYBR Green kit (ThermoFisher) to detect LACV. The following primers were used: forward (ATTCTACCCGCTGACCATTG) and reverse (GTGAGAGTGCCATAGCGTTG) (15). qPCR was performed using a QuantStudio 3Flex instrument (Applied Biosystems). QuantStudio Real- Time PCR software was used to extract data. GAPDH was used as a control of expression for qPCR samples.

### Infection of mice with LACV and timepoint collection

All animal studies were approved by the Rutgers University IACUC (protocol number 202300007) and performed in accordance with protocols set forth by the National Institutes of Health *Guide for the Care and Use of Laboratory Animals* and ARRIVE guidelines. Wildtype (C57BL/6) mice were purchased from Jackson Laboratories and maintained in a breeding colony at the Child Health Institute of New Jersey. Weanling (3 weeks old) and adult mice (8 weeks old) were inoculated with 10^3^ PFU of LACV in phosphate-buffered saline (PBS) intraperitoneally in a volume of 100 μl/mouse. Mice were observed daily for signs of neurological disease. Animals with clinical signs were scored and humanely euthanized if they demonstrated clear signs of neurological disease, including seizures plus lethargy. For the clinical scoring graphs, the percentage of mice with neurological disease was calculated based on the number of weanling or adult mice out of the total number of surviving animals for each age groups. Mice were sacrificed and brains were collected at the following timepoints: 2, 3, 4, 5, and 7 days post-infection (dpi) for weanlings and 4, 5, 6, 7, 10, and 20 dpi for adults and were stored in RNAse away, as previously described (20, 22). Viral RNA was extracted as described.

### Rapid test strip image analysis and viral quantification

To estimate viral loads from rapid test strip images, we first established a standard curve using known viral RNA copies. LACV RNA was serially diluted to generate concentrations ranging from 10^6^ to 0 copies for RT-RPA testing with results visualization via LFA strips. Resulting test line bands were imaged using under consistent lighting conditions. Image processing and pixel intensity quantification were performed using ImageJ software (NIH) (27, 42–44). The pixel intensity of the test line for each dilution was recorded and plotted against the corresponding viral copy number to generate a standard curve. A linear regression model was fitted to the standard curve, and the equation y = 10^(m*x+b)^ was derived, where x represents the pixel intensity, and y corresponds to the estimated viral copy number. This equation was used to interpolate viral loads for test samples based on their measured pixel intensities.

The test line intensity for each weanling and adult mouse sample was measured using ImageJ and converted to viral copies using the standard curve equation. Pixel intensities for adult and weanling samples were plotted on the standard curve to estimate viral load dynamics over time. Graphical representations were performed using Python (Matplotlib, NumPy, SciPy, and Seaborn). The standard curve was plotted as a black straight live, generated using linear repression.

## Results

### Screening of RT-RPA primer pairs for LACV detection

To identify a genome segment best suited for specific and sensitive detection of LACV, we performed a percentage identity matrix analysis on the S, M, and L segments from a panel of CSG viruses. These included LACV, SSHV, Khatanga virus, California Encephalitis virus, San Angelo virus, Lumbo virus, and Tahnya virus, and one virus from the Melaka virus family, Jamestown Canyon virus. Based on homology analyses of DNA sequences, the S, M, and L segments displayed homology ranging from 76.1-88%, 69- 78.8%, and 73.7-81.6%, respectively (Fig. 1, Table 1). Since the M segment exhibited the greatest divergence between LACV and other CSG viruses, we selected it as the target for LACV-specific primer design.

**Fig. 1.**
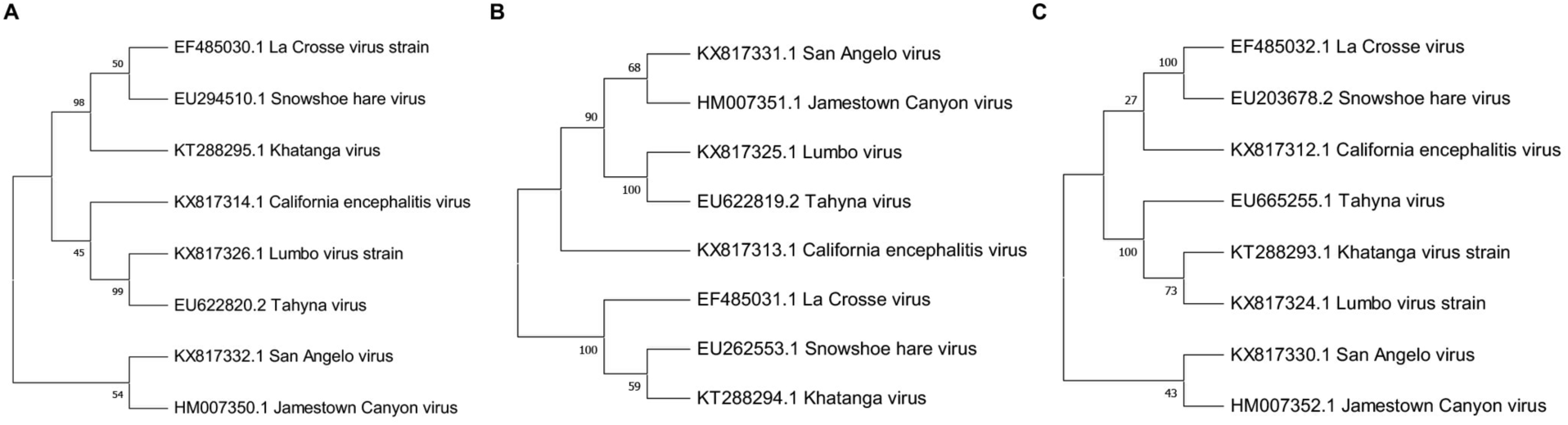
Phylogenetic trees of the S, M, L segments of LACV and related bunyaviruses. Phylogenetic trees were constructed from aligning nucleotide sequences of the A) small (S), B) medium (M), and C) large (L) segments from bunyaviruses within the CSG, including La Crosse virus, Snowshoe hare virus, Khatanga virus, California Encephalitis virus, San Angelo virus, Lumbo virus, and Tahnya virus, and Melaka virus (MELV) family, including Jamestown Canyon virus. The sequences were aligned using Clustal omega, and the resulting alignments were used to construct maximum-likelihood trees based on the Tamura-Nei model and 500 bootstrapping replications using Mega11. Associated accession numbers are included by the name of each virus for each tree.

**Table 1.** Nucleic acid sequence homology of the S, M, and L segments between LACV and other related bunyaviruses.

| Virus | S Segment Homology | M Segment Homology | L Segment Homology |
| --- | --- | --- | --- |
| La Crosse virus | 100% | 100% | 100% |
| Snowshoe hare virus | 88% | 78.8% | 81.6% |
| Khatanga virus | 87% | 77.5% | 78% |
| California encephalitis virus | 79.8% | 73.6% | 76.9% |
| Tahyna virus | 82.2% | 73% | 77.3% |
| Lumbo virus | 80.7% | 72.3% | 76.7% |
| San Angelo virus | 80.1% | 70.5% | 77.1% |
| Jamestown Canyon virus | 76.1% | 69% | 73.7% |

Four primer pairs were generated, each producing amplicons of at least 100 base pairs. These primers were tested at 37°C, 39°C, and 42°C for 20 minutes. Primer pair 3 at 39°C yielded the clearest and most intense amplification band (208 base pairs [bp]) and was chosen for subsequent optimization (Fig. 2). Primer pair 4 also showed reliable results (240 bp), but was slightly less optimal. Notably, the RT-RPA amplified products were analyzed via agarose gel electrophoresis without prior purification, enabling efficient screening of primer pairs.

**Fig. 2.**
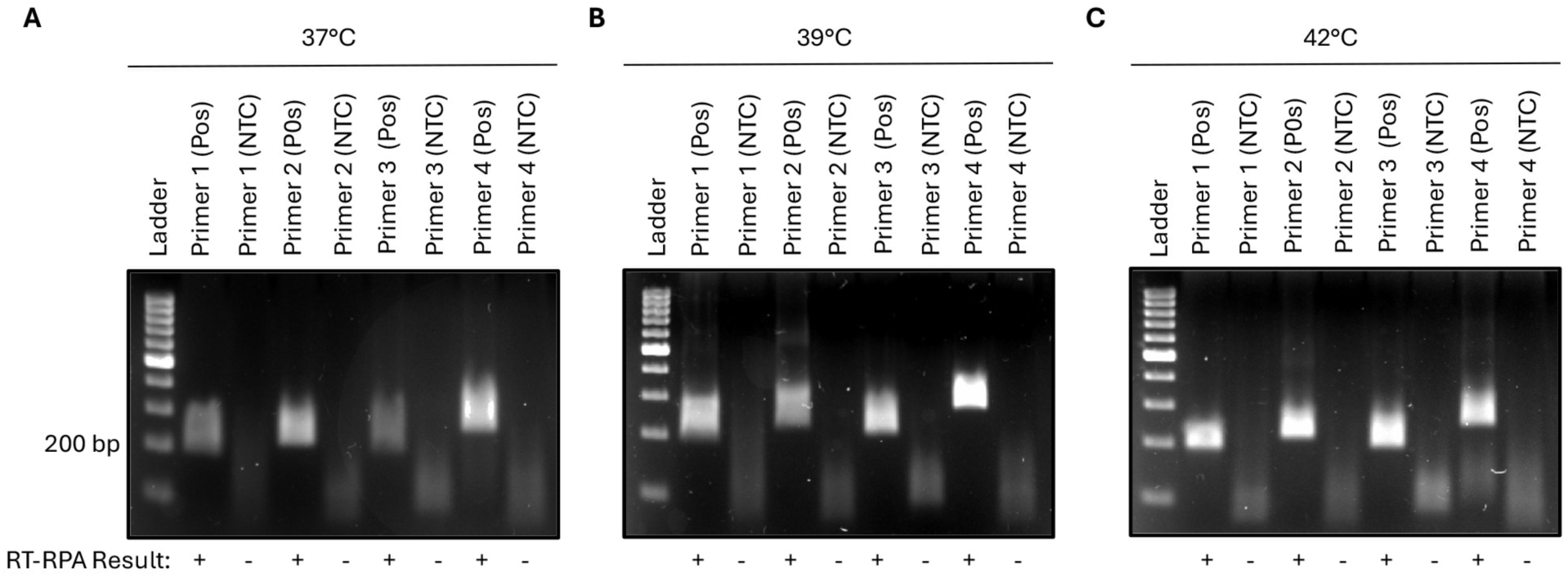
Primer screen RT-RPA testing. Primer screen RT-RPA testing at A) 37°C, B) 39°C, and C) 42°C using 4 different primer pairs. Primer pair 3 at 39°C was chosen for further experimentation. The RT-RPA amplified products were not purified prior to running on the agarose gel. Pos, positive control, La crosse virus cDNA input. NTC, no template control, water input. +, positive result. -, negative result.

### Assessing LACV RT-RPA amplification time and analytical sensitivity and specificity via gel electrophoresis

After identifying the best performing primer pair, we conducted a time course study to determine the minimal incubation period necessary for effective RT-RPA detection of LACV. Reactions were incubated for 5, 10, 20, and 60 minutes before the amplicons were purified and visualized by agarose gel electrophoresis. Although faint bands appeared as early as 10 minutes, we observed optimal band clarity and intensity at 20 minutes (Fig. 3A).

**Fig. 3.**
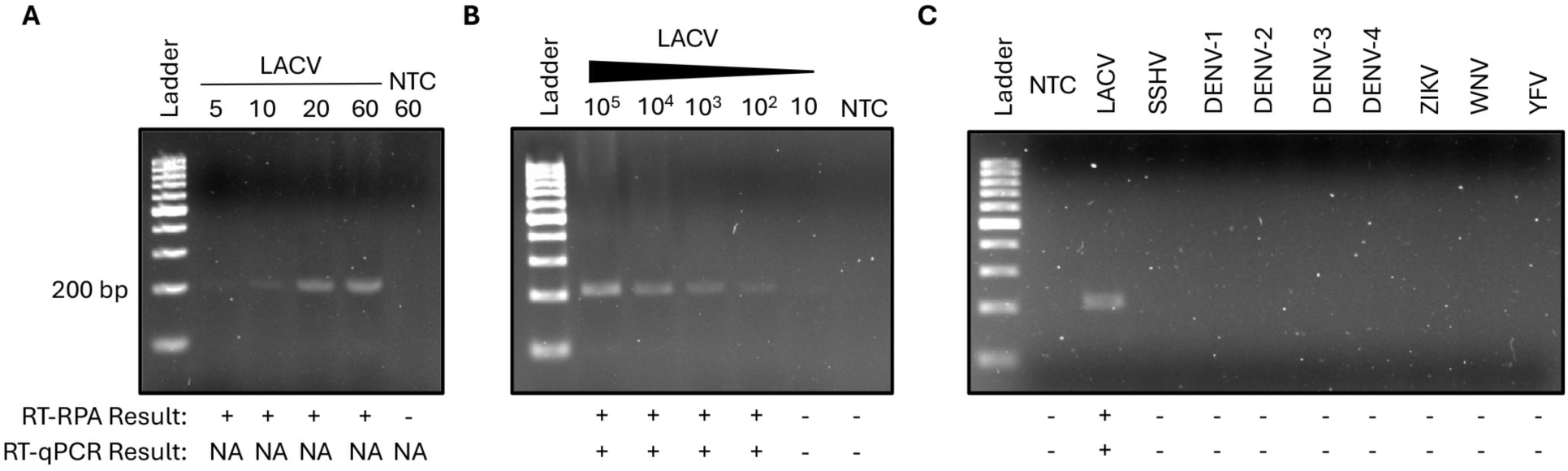
Time Course, limit of detection, and cross-reactivity RT-RPA testing with visualization of results via gel electrophoresis. A) Time course tests were performed at 5 minutes, 10 minutes, 20 minutes and 60 minutes. The time course was optimized at 20 minutes. RT-qPCR testing in the time course experiment was not applicable (NA). B) Viral RNA from 10^5^ copies to 0 copies was reverse transcribed to cDNA and used in RT-RPA and RT-qPCR testing. The RT-RPA limit of detection was determined to be 100 viral copies. C) RT-RPA was performed using cDNA from La Crosse virus (LACV), Snowshoe hare virus (SSHV), dengue virus serotype 1 (DENV-1), dengue virus serotypes 2 (DENV-2), dengue virus serotype 3 (DENV-3), dengue virus serotype 4 (DENV-4), Zika virus (ZIKV), West Nile virus (WNV), or yellow fever virus (YFV). No cross-reactivity was observed. The RT-RPA products were purified prior to running on the agarose gel. NTC, no template control, water input. +, positive result. -, negative result.

Next, we evaluated the RT-RPA assay’s analytical sensitivity using a ten-fold serial dilution of LACV RNA, ranging from 10^5^ to 0 input copies in the RT step. The RT-RPA reactions were purified and then assessed via agarose gel electrophoresis. The LOD was established at 100 copies (Fig. 3B), similar to the RT-qPCR assay. To verify the RT-RPA assay’s specificity, we tested RNA extracted from other arboviruses, including SSHV, dengue virus serotypes 1-4 (DENV-1-4), Zika virus (ZIKV), West Nile virus (WNV), and yellow fever virus (YFV). Similarly, the RT-RPA reactions were purified and then assessed via agarose gel electrophoresis. No RT-RPA or RT-qPCR amplification occurred in any non- LACV samples, confirming the specificity of our primer pair (Fig. 3C).

### Integration of the LACV RT-RPA assay with an LFA

To enable the potential point-of-care diagnosis of LACV, a lateral flow assay (LFA) was developed and integrated with the LACV RT-RPA assay. We employed the same unlabeled forward primer, but the reverse primer was adapted to contain a 5’-biotin tag, combined with a short hybridization step in which the probe was labeled with a 5’-FAM tag. After RT-RPA and hybridization, the product was exposed to the LFA strip, containing detection lines coated with streptavidin on the test area and anti-rabbit antibody on the control area. Gold nanoparticle-conjugated anti-rabbit-FITC antibodies generated the visual signal upon binding to hybridization product containing the FAM and biotin labels.

The LFA exhibited the same LOD of 100 copies (Fig. 4A). Similarly, the LFA also did not cross-react with SSHV and the other geographically relevant arboviruses (Fig. 4B). These results confirm the LFA’s utility as a potential rapid and field-deployable tool for LACV detection.

**Fig. 4.**
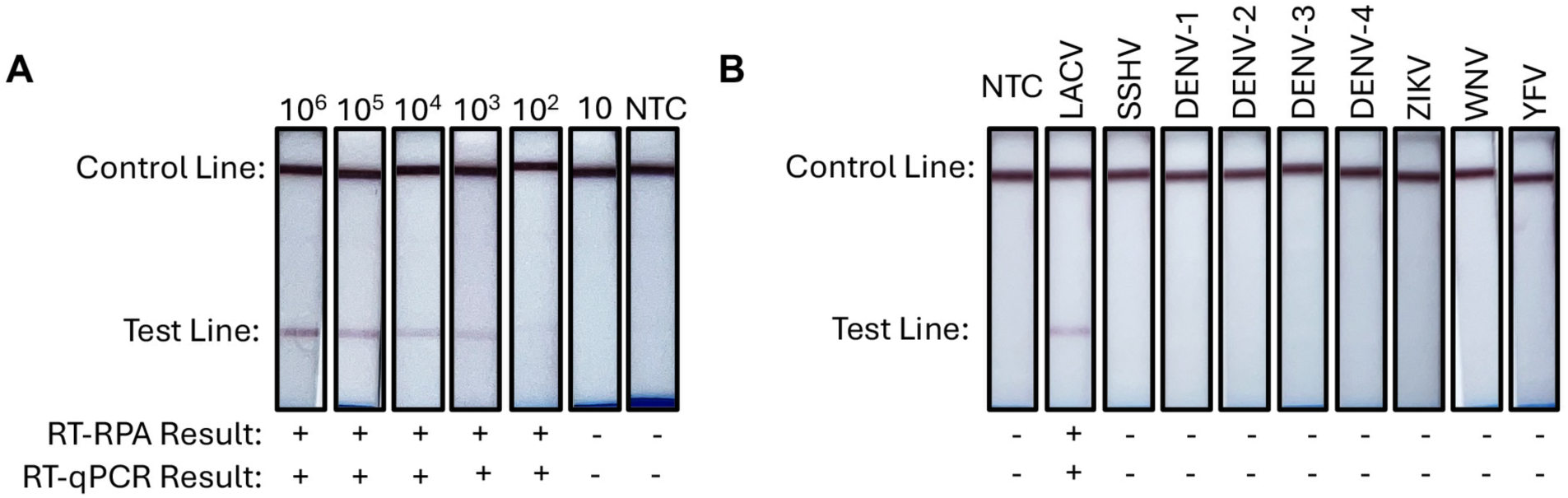
RT-RPA limit of detection and cross-reactivity testing with visualization of results via LFA strips. A) Viral RNA from 10^6^ copies to 0 copies were reverse transcribed to cDNA and used in RT-RPA and RT-qPCR testing. The limit of detection was determined to be 100 viral copies. B) RT-RPA and RT-qPCR testing was performed using cDNA from La Crosse virus (LACV), Snowshoe hare virus (SSHV), dengue virus serotype 1 (DENV-1), dengue virus serotypes 2 (DENV-2), dengue virus serotype 3 (DENV-3), dengue virus serotype 4 (DENV-4), Zika virus (ZIKV), West Nile virus (WNV), or yellow fever virus (YFV). No cross-reactivity was observed with either RT-RPA or RT-qPCR testing. NTC, no template control, water input. +, positive result. -, negative result.

### Evaluation of age-dependent detection and clearance of LACV in weanling and adult mouse brains

To probe how host age modulates LACV neuroinvasion, we utilized both weanling and adult C57BL/6 mice. Animals were inoculated intraperitoneally with 10^3^ plaque forming units (PFU) of LACV, and brains were collected at distinct dpi. Weanling mice were sacrificed and sampled at 2, 3, 4, 5, and 7 dpi, whereas adult mice were sacrificed and sampled at 4, 5, 6, 7, 10, and 20 dpi (Supplementary Fig. 1A). RNA from brains was extracted, reverse transcribed to cDNA, and evaluated using RT-qPCR as well as the newly developed RT-RPA test, with results visualization by LFA strips and agarose gel electrophoresis.

We observed that LACV accessed the brains of weanling mice as early as 4 dpi and was detectable in all mice at the 5-7 dpi timepoints (Fig. 5A, Supplementary Fig. 1B). The onset of symptoms coincided with LACV access to the CNS, with abnormal gait observed in 11% (one out of nine remaining weanling mice at 4 dpi), and 100% of the remaining weanlings exhibiting severe neurological symptoms between 5-7 dpi (Fig. 5B). In adult mice, LACV was first detected at 5 dpi, plateauing between 6-10 dpi (Fig. 5C, Supplementary Fig. 1C). By 20 dpi, LACV was undetectable in the surviving adults, suggesting robust viral clearance. In contrast to weanlings, 100% of adults exhibited no symptoms between 4-7 dpi, with abnormal gait observed in 16.7% (one out of six remaining adult mice) at 10 dpi (Fig. 5D). The 20 dpi timepoint included only two mice as one mouse in the 20 dpi cohort succumbed to the infection at 12 dpi and could not be included in the analysis.

**Fig. 5.**
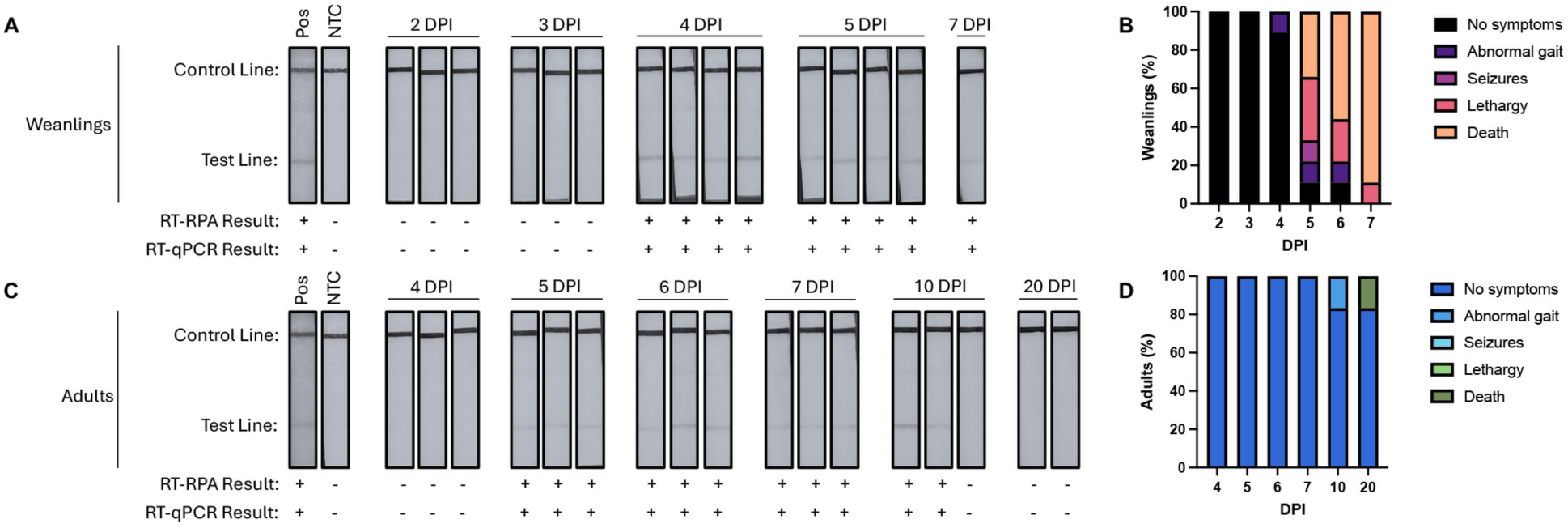
LACV neuroinvasion into the brain of weanling and adult mice visualized by LFA strips. Mice were infected with 1000 plaque forming units of LACV and were humanely sacrificed and brains were collected 2-7 days post-infection (DPI) for weanlings and 4-20 DPI for adults. A) LACV was detected in the brain in weanling mice by both RT-RPA and RT-qPCR as early as 4 DPI. B) Bar chart representing percent weanling mice found with neurological symptoms. There was an onset of clinical symptoms by 4 DPI in weanlings, with worsening of neurological symptoms between 5-7 DPI. C) LACV was detected in the brain in adult mice by both RT-RPA and RT-qPCR by 5 DPI with clearance of the virus by 20 DPI. D) Bar chart representing percent adult mice found with neurological symptoms. There were no symptoms exhibited in adults between 4-7 DPI; however, one adult exhibited neurological symptom by 12 DPI and could not be included in the analysis. Two remaining adults were asymptomatic throughout the course of the study. Pos, positive control, La crosse virus cDNA input. NTC, no template control, water input. +, positive result. -, negative result.

Furthermore, viral load quantification was derived from a standard curve generated from known LACV RNA copies (Fig. 4A). The standard curve demonstrated a strong correlation between pixel intensity and viral copies (R^2^ > 0.99), enabling estimation of LACV burden in the brain. Estimated viral copies in weanling brains at peak infection (4-5 dpi) ranged from ∼10^4.5^ to ∼10^6^, consistent with high neuroinvasive potential at early time points (Fig. 6A-B). Peak viral burden in adult brains was notably lower than in weanlings between 5-10 dpi, with estimated viral loads between ∼10^2^ to ∼10^3^ copies between (Fig. 6A-B). However, there was one adult with high viral load at 10 dpi (Fig. 6A-B).

**Fig. 6.**
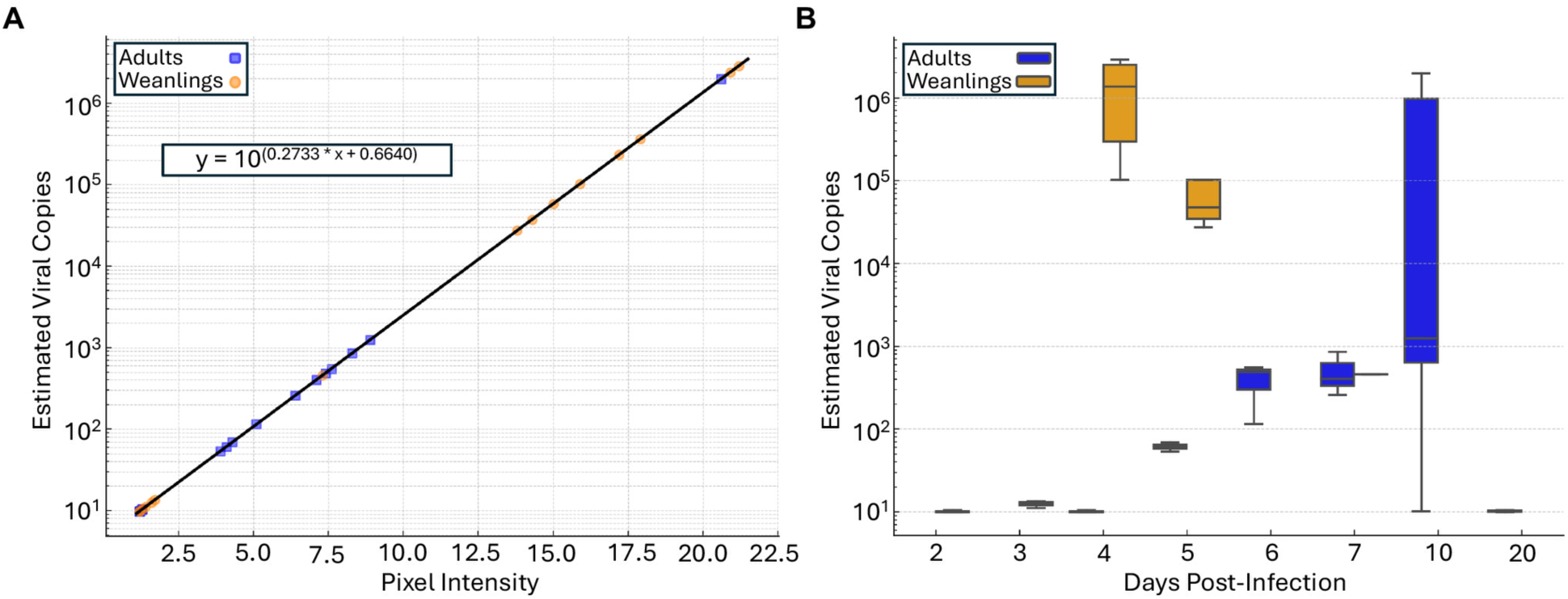
Estimation of LACV viral burden in the brain from LFA strips. A) Test line pixel intensities for LACV-infected weanling (orange circles) and adult (blue squares) mice were plotted on the standard curve to estimate viral load dynamics over time. The standard curve is plotted as a black straight line, generated using linear regression (equation of the standard curve is included in the box). A logarithmic scale was used for viral copies to better represent dynamic changes in viral loads across multiple orders of magnitude. B) Box-and-whisker plot represents the distribution of estimated viral copies in weanlings (orange) and adults (blue) at each time point.

## Discussion

In this study, we developed, optimized, and validated an RT-RPA assay to detect LACV. Traditional RT-qPCR techniques, although highly sensitive, rely on costly equipment and specialized laboratory settings, and often lose sensitivity as viral titers decline (24, 25). Serological tools, meanwhile, necessitate confirmatory plaque reduction neutralization tests (PRNTs) to differentiate among closely related arboviruses—a process that can be time-intensive and cumbersome for large-scale or field-based testing (26, 45). Our RT-RPA assay addresses these limitations by demonstrating efficient and robust LACV detection with a LOD of 100 RNA copies and without cross-reaction. This performance was replicated in both agarose gel electrophoresis and LFA strips, underscoring the assay’s versatility and potential utility in resource-constrained environments where on-site diagnostics could significantly improve patient management and epidemiological surveillance.

An important aspect of our assay’s development was the primer design targeting the M segment. Using bioinformatic analysis of multiple CSG viruses, we observed greater sequence divergence in the M segment between LACV and closely related CSG viruses. By designing primers and a probe that map specifically to these divergent regions, we achieved high specificity without any cross-reactivity against SSHV and other geographically relevant arboviruses (46). These findings confirm that our assay is well suited to environments where multiple arboviruses circulate simultaneously. Moreover, the integrated approach of utilizing an unpurified RT-RPA product for hybridization plus results readout via an LFA strip reliably produced clear test lines, suggesting a potentially straightforward workflow for use in clinics or field settings lacking sophisticated equipment. In future iterations, combining the RT and RPA steps into a single reaction may further streamline testing by potentially eliminating the need for RNA extraction and cDNA synthesis.

Beyond its diagnostic innovation, our study provides important new insights into LACV pathogenesis, particularly the impact of host age on the temporal neuroinvasion dynamics. While the progression of LACV neuroinvasion in weanling mice has been well studied, the dynamics in adult mice remained poorly characterized, and a direct comparative analysis between weanlings and adults had not been previously investigated (39). Using a serial euthanasia approach, we assessed LACV detection in the brain of both weanling and adult mice at predefined intervals (2–7 dpi for weanlings; 4–20 dpi for adults). We demonstrated that weanling mice exhibited detectable virus in the brain as early as 4 dpi, with sustained high viral loads between 5-7 dpi, and most weanling mice displaying severe CNS pathology by the end of the study window. Conversely, in adult mice, LACV was first detected in the brain at 5 dpi—indicating a delayed onset of neuroinvasion compared to weanlings—and exhibited plateauing levels between 6–10 dpi. Notably, some adults still harbored detectable virus at 10 dpi, yet all surviving adults cleared the virus within the brain completely by 20 dpi.

While survival outcomes were not directly assessed in this study due to our serial euthanasia design, extensive prior research has consistently demonstrated that weanlings typically succumb by 7 dpi, whereas adult mice exhibit minimal to no mortality (20–22, 39, 40). However, our estimation of LACV burden based on LFA strip image analysis demonstrated notably higher viral loads in the brains of weanlings compared to adults. Together with the current study, these findings suggest that adult mice experience a slower initial spread of virus into the CNS relative to weanlings, potentially allowing more time for immune responses to mitigate viral replication and promote survival. Although further immunological assays would be required to pinpoint the specific immune responses involved, the delayed neuroinvasion and ultimately effective viral clearance in adults likely underlie their more favorable clinical outcomes compared to younger hosts (21). Our recent analysis of T cell immunity demonstrates significant disparities in the cellular response to LACV in weanling and adult mice (47). Adult mice mount significantly more polyfunctional CD4^+^ and CD8^+^ T cell responses compared to minimal responses in weanlings (47).

Despite the strengths of our study, several limitations should be acknowledged. First, while our RT-RPA assay demonstrated high sensitivity and specificity for LACV detection, we did not evaluate its performance using human clinical samples. Given the rarity of confirmed LACV cases and the challenges in obtaining clinical specimens, validation in patient samples remains an important next step. Additionally, our study relied on in vitro and murine models to assess viral detection and neuroinvasion. While these models provide valuable insights, human pathophysiology may differ, necessitating further validation in human-derived tissues or clinical studies. Another limitation is that while our assay successfully distinguished LACV from closely related CSG viruses, broader validation against a wider panel of co-circulating arboviruses would further strengthen its diagnostic utility.

Furthermore, a potential limitation of RT-RPA/LFA is reduced sensitivity at early and late stages of infection, when viral titers are low. While our assay demonstrated a limit of detection of 100 copies, further optimization is needed to enhance sensitivity for clinical applications. Future improvements could include incorporating a pre-amplification step to increase RNA yield, using fluorescent or nanoparticle-based lateral flow detection to enhance signal amplification, and developing a multiplexed RT-RPA approach targeting multiple genomic regions to improve detection across a range of viral loads. Finally, future work should explore the integration of RT and RPA into a single-tube reaction to further simplify the assay workflow, enhancing its feasibility for field deployment.

## Conclusion

Taken together, our work underscores a dual contribution to the field: we introduce a highly sensitive and specific RT-RPA assay for LACV detection, and we leverage this assay to enhance our understanding of age-dependent neuroinvasion in a murine model.

Importantly, the RT-RPA method is amenable to further refinements, such as one-step RT-RPA protocols or improved probe chemistries, which could streamline its use under outbreak conditions. From a public health perspective, quicker diagnoses in endemic or high-risk regions would facilitate timely interventions, especially for pediatric populations at greatest risk of severe outcomes. Meanwhile, our findings on the timing and efficiency of viral invasion in adult versus weanling brains point toward an inherent advantage conferred by age-related immune maturity, reinforcing the notion that prophylactic or early therapeutic measures could be particularly impactful in children. Overall, these advancements promise to mitigate the burden of LACV disease and enrich our broader comprehension of arboviral neuroinvasion.

## Ethics approval and consent to participate

Not applicable.

## Consent for publication

Not applicable.

## Availability of data and material

All data generated or analyzed during this study are included in the published article and its supplementary information files.

## Author Contributions

LL designed probes and primers, planned, ran most of the experiments described above, analyzed experimental data, and wrote the original draft of manuscript. RA assisted with mouse infections, mouse monitoring, and brain collection. SS helped with the development of primers, probes, and protocol optimization. PBT helped with RNA extractions. BBH developed the ideas for the project, as well as supervised the work, and planned and carried out experiments. All authors contributed to the interpretation of the data and played a role in reviewing and editing the manuscript.

## Acknowledgements

We would like to thank Rutgers Global Health Institute, Rutgers Robert Wood Johnson Medical School, and The Child Health Institute of New Jersey for their continued support.

---

**Supplementary Fig. 1.**
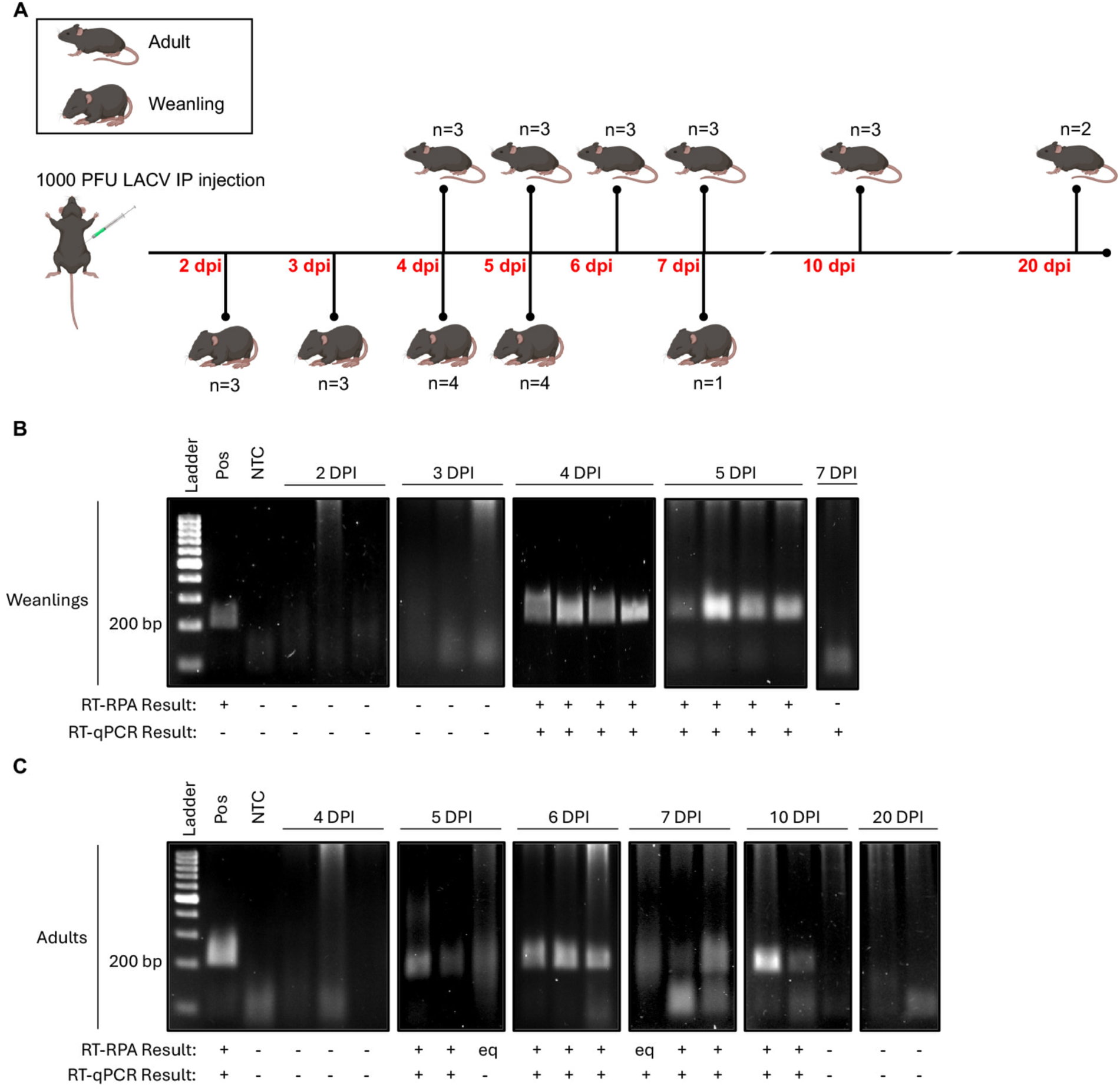
Serial euthanasia schematic and LACV neuroinvasion into the brain of weanling and adult mice visualized by gel electrophoresis. A) Mice were infected with 1000 plaque forming units of LACV and mice were humanely sacrificed, and brains were collected 2-7 days post-infection (dpi) for weanlings and 4-20 dpi for adults. B) LACV was detected in the brain in weanling mice by both RT-RPA and RT-qPCR as early as 4 DPI. The RT-RPA amplified products were not purified prior to running on the agarose gel. C) LACV was detected in the brain in adult mice by both RT-RPA and RT-qPCR by 5 DPI with clearance of the virus by 20 DPI. The RT-RPA amplified products were not purified prior to running on the agarose gel. Pos, positive control, La crosse virus cDNA input. NTC, no template control, water input. +, positive result. -, negative result. eq, equivocal result.

**Supplementary Table 1.**
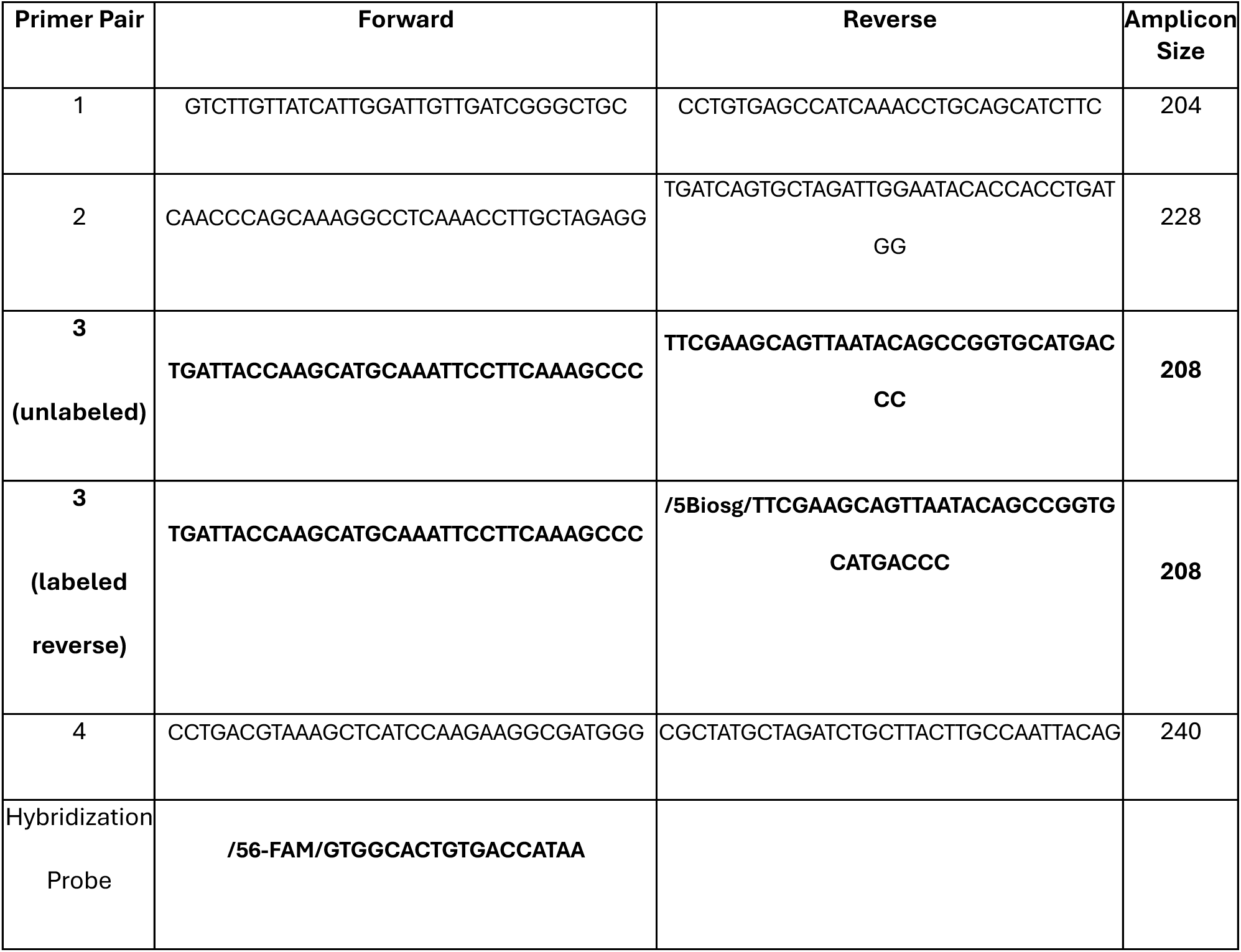
RT-RPA primer pairs and hybridization probe.

## References

1. Alatrash R, Herrera BB. The Adaptive Immune Response against Bunyavirales. Viruses. 2024;16(3).

2. Evans AB, Peterson KE. Throw out the Map: Neuropathogenesis of the Globally Expanding California Serogroup of Orthobunyaviruses. Viruses. 2019;11(9).

3. Soldan SS, Plassmeyer ML, Matukonis MK, Gonzalez-Scarano F. La Crosse virus nonstructural protein NSs counteracts the effects of short interfering RNA. J Virol. 2005;79(1):234–44.

4. Blakqori G, Delhaye S, Habjan M, Blair CD, Sanchez-Vargas I, Olson KE, et al. La Crosse bunyavirus nonstructural protein NSs serves to suppress the type I interferon system of mammalian hosts. J Virol. 2007;81(10):4991–9.

5. Soldan SS, Hollidge BS, Wagner V, Weber F, Gonzalez-Scarano F. La Crosse virus (LACV) Gc fusion peptide mutants have impaired growth and fusion phenotypes, but remain neurotoxic. Virology. 2010;404(2):139–47.

6. Reese SM, Blitvich BJ, Blair CD, Geske D, Beaty BJ, Black WCt. Potential for La Crosse virus segment reassortment in nature. Virol J. 2008;5:164.

7. Bennett RS, Ton DR, Hanson CT, Murphy BR, Whitehead SS. Genome sequence analysis of La Crosse virus and in vitro and in vivo phenotypes. Virol J. 2007;4:41.

8. Arragain B, Durieux Trouilleton Q, Baudin F, Provaznik J, Azevedo N, Cusack S, et al. Structural snapshots of La Crosse virus polymerase reveal the mechanisms underlying Peribunyaviridae replication and transcription. Nat Commun. 2022;13(1):902.

9. Day CA, Odoi A, Trout Fryxell R. Geographically persistent clusters of La Crosse virus disease in the Appalachian region of the United States from 2003 to 2021. PLoS Negl Trop Dis. 2023;17(1):e0011065.

10. McJunkin JE, Khan RR, Tsai TF. California-La Crosse encephalitis. Infect Dis Clin North Am. 1998;12(1):83–93.

11. Goldman T, Hamer DH. Current Status of La Crosse Virus in North America and Potential for Future Spread. Am J Trop Med Hyg. 2024;110(5):850–5.

12. Haddow AD, Odoi A. The incidence risk, clustering, and clinical presentation of La Crosse virus infections in the eastern United States, 2003-2007. PLoS One. 2009;4(7):e6145.

13. CDC. La Cross virus [Available from: https://www.cdc.gov/la-crosse-encephalitis/about/index.html].

14. Gaensbauer JT, Lindsey NP, Messacar K, Staples JE, Fischer M. Neuroinvasive arboviral disease in the United States: 2003 to 2012. Pediatrics. 2014;134(3):e642–50.

15. Evans AB, Winkler CW, Peterson KE. Differences in Neuropathogenesis of Encephalitic California Serogroup Viruses. Emerg Infect Dis. 2019;25(4):728–38.

16. Lau L, Wudel B, Kadkhoda K, Keynan Y. Snowshoe Hare Virus Causing Meningoencephalitis in a Young Adult From Northern Manitoba, Canada. Open Forum Infect Dis. 2017;4(3):ofx150.

17. Boutzoukas AE, Freedman DA, Koterba C, Hunt GW, Mack K, Cass J, et al. La Crosse Virus Neuroinvasive Disease in Children: A Contemporary Analysis of Clinical/Neurobehavioral Outcomes and Predictors of Disease Severity. Clin Infect Dis. 2023;76(3):e1114–e22.

18. Harding S, Greig J, Mascarenhas M, Young I, Waddell LA. La Crosse virus: a scoping review of the global evidence. Epidemiol Infect. 2018;147:e66.

19. Day CA, Byrd BD, Trout Fryxell RT. La Crosse virus neuroinvasive disease: the kids are not alright. J Med Entomol. 2023;60(6):1165–82.

20. Basu R, Ganesan S, Winkler CW, Anzick SL, Martens C, Peterson KE, et al. Identification of age-specific gene regulators of La Crosse virus neuroinvasion and pathogenesis. Nat Commun. 2023;14(1):2836.

21. Alatrash R, Vaidya V, Herrera BB. Age-specific dynamics of neutralizing antibodies, cytokines, and chemokines in response to La Crosse virus infection in mice. J Virol. 2024;98(12):e0176224.

22. Basu R, Nair V, Winkler CW, Woods TA, Fraser IDC, Peterson KE. Age influences susceptibility of brain capillary endothelial cells to La Crosse virus infection and cell death. J Neuroinflammation. 2021;18(1):125.

23. Teleron AL, Rose BK, Williams DM, Kemper SE, McJunkin JE. La Crosse Encephalitis: An Adult Case Series. Am J Med. 2016;129(8):881–4.

24. Wasieloski LP, Jr., Rayms-Keller A, Curtis LA, Blair CD, Beaty BJ. Reverse transcription-PCR detection of LaCrosse virus in mosquitoes and comparison with enzyme immunoassay and virus isolation. J Clin Microbiol. 1994;32(9):2076–80.

25. Watzinger F, Ebner K, Lion T. Detection and monitoring of virus infections by real- time PCR. Mol Aspects Med. 2006;27(2-3):254–98.

26. Lambert AJ, Nasci RS, Cropp BC, Martin DA, Rose BC, Russell BJ, et al. Nucleic acid amplification assays for detection of La Crosse virus RNA. J Clin Microbiol. 2005;43(4):1885–9.

27. Salcedo N, Sena BF, Qu X, Herrera BB. Comparative Evaluation of Rapid Isothermal Amplification and Antigen Assays for Screening Testing of SARS-CoV-2. Viruses. 2022;14(3).

28. Evans AB, Peterson KE. Cross reactivity of neutralizing antibodies to the encephalitic California Serogroup orthobunyaviruses varies by virus and genetic relatedness. Sci Rep. 2021;11(1):16424.

29. Daher RK, Stewart G, Boissinot M, Bergeron MG. Recombinase Polymerase Amplification for Diagnostic Applications. Clin Chem. 2016;62(7):947–58.

30. Faye M, Abd El Wahed A, Faye O, Kissenkotter J, Hoffmann B, Sall AA, et al. A recombinase polymerase amplification assay for rapid detection of rabies virus. Sci Rep. 2021;11(1):3131.

31. Vasileva Wand NI, Bonney LC, Watson RJ, Graham V, Hewson R. Point-of-care diagnostic assay for the detection of Zika virus using the recombinase polymerase amplification method. J Gen Virol. 2018;99(8):1012–26.

32. Lobato IM, O’Sullivan CK. Recombinase polymerase amplification: Basics, applications and recent advances. Trends Analyt Chem. 2018;98:19–35.

33. Sainetra Sridhar PBT, Lily Lumkong, Eduardo Martins Netto, Carlos Brites, Wei-Kung Wang C, Bobby Brooke Herrera. RT-RPA as a dual tool for detection and phylogenetic analysis of epidemic arthritogenic alphaviruses. Scientific Reports. 2024;14.

34. Jauset-Rubio M, Svobodova M, Mairal T, McNeil C, Keegan N, Saeed A, et al. Ultrasensitive, rapid and inexpensive detection of DNA using paper based lateral flow assay. Sci Rep. 2016;6:37732.

35. Shelite TR, Uscanga-Palomeque AC, Castellanos-Gonzalez A, Melby PC, Travi BL. Isothermal recombinase polymerase amplification-lateral flow detection of SARS-CoV-2, the etiological agent of COVID-19. J Virol Methods. 2021;296:114227.

36. Ghosh DK, Kokane SB, Gowda S. Development of a reverse transcription recombinase polymerase based isothermal amplification coupled with lateral flow immunochromatographic assay (CTV-RT-RPA-LFICA) for rapid detection of Citrus tristeza virus. Sci Rep. 2020;10(1):20593.

37. Davis J, Atkins C, Doyle M, Williams C, Boyce R, Byrd B. Endemic La Crosse Virus Neuroinvasive Disease in North Carolina Residents: 2000-2020. N C Med J. 2024;85(4):289–95.

38. Taylor KG, Woods TA, Winkler CW, Carmody AB, Peterson KE. Age-dependent myeloid dendritic cell responses mediate resistance to la crosse virus-induced neurological disease. J Virol. 2014;88(19):11070–9.

39. Bennett RS, Cress CM, Ward JM, Firestone CY, Murphy BR, Whitehead SS. La Crosse virus infectivity, pathogenesis, and immunogenicity in mice and monkeys. Virol J. 2008;5:25.

40. Evans AB, Winkler CW, Peterson KE. Differences in neuroinvasion and protective innate immune pathways between encephalitic California Serogroup orthobunyaviruses. PLoS Pathog. 2022;18(3):e1010384.

41. Qian J, Boswell SA, Chidley C, Lu ZX, Pettit ME, Gaudio BL, et al. An enhanced isothermal amplification assay for viral detection. Nat Commun. 2020;11(1):5920.

42. Harmon A, Chang C, Salcedo N, Sena B, Herrera BB, Bosch I, et al. Validation of an At-Home Direct Antigen Rapid Test for COVID-19. JAMA Netw Open. 2021;4(8):e2126931.

43. Salcedo N, Harmon A, Herrera BB. Pooling of Samples for SARS-CoV-2 Detection Using a Rapid Antigen Test. Front Trop Dis. 2021;2:707865.

44. Salcedo N, Reddy A, Gomez AR, Bosch I, Herrera BB. Monoclonal antibody pairs against SARS-CoV-2 for rapid antigen test development. PLoS Negl Trop Dis. 2022;16(3):e0010311.

45. Calisher CH, Pretzman CI, Muth DJ, Parsons MA, Peterson ED. Serodiagnosis of La Crosse virus infections in humans by detection of immunoglobulin M class antibodies. J Clin Microbiol. 1986;23(4):667-71.

46. Pierson TC, Diamond MS. The continued threat of emerging flaviviruses. Nat Microbiol. 2020;5(6):796–812.

47. Alatrash R, Herrera BB. Immunodominant structural proteins Gc and N drive T cell-mediated protection against La Crosse virus. bioRxiv. 2025:2025.02. 25.640063.

